# Microfluidic bioprinting for the in vitro generation of novel biomimetic human testicular tissues

**DOI:** 10.1101/2021.06.04.447126

**Authors:** Meghan Robinson, Erin Bedford, Luke Witherspoon, Stephanie M. Willerth, Ryan Flannigan

**Affiliations:** Vancouver Prostate Centre, 2660 Oak St, Vancouver, British Columbia, Canada V6H 3Z6; Aspect Biosystems, 1781 W 75th Ave, Vancouver, British Columbia, Canada V6P 6P2; Department of Urologic Sciences, University of British Columbia, 2329 West Mall, Vancouver, British Columbia, Canada V6T 1Z4; Department of Urology, The Ottawa Hospital, 501 Smyth Rd, Ottawa, Ontario, Canada K1H 8L6; Division of Medical Sciences, University of Victoria, 3800 Finnerty Road, Victoria, British Columbia, Canada V8P 5C2; Department of Mechanical Engineering, University of Victoria, Victoria, British Columbia, Canada; School of Biomedical Engineering, University of British Columbia, 2329 West Mall, Vancouver, British Columbia, Canada V6T 1Z4; Department of Urology, Weill Cornell Medicine, 1300 York Ave, New York, New York 10065, United State

**Author notes:** **Corresponding author** Ryan Flannigan, MD Department of Urologic Sciences, University of British Columbia, Vancouver, British Columbia Gordon & Leslie Diamond Health Care Centre Level 6, 2775 Laurel Street Vancouver, BC Canada V5Z 1M9 Phone number.

**Keywords:** Bioprinting, spermatogenesis, seminiferous tubules, fertility, biomimetics, coaxial

## Abstract

Advances in cancer treatments have greatly improved pediatric cancer survival rates, leading to quality of life considerations and in particular fertility restoration. Accordingly, pre-pubertal patients have the option to cryopreserve testicular tissue for experimental restorative therapies, including *in vitro* spermatogenesis, wherein testicular tissue is engineered *in vitro* and spermatozoa are collected for *in vitro* fertilization (IVF). Current *in vitro* systems have been unable to reliably support the generation of spermatozoa from human testicular tissues, likely due to the inability for the dissociated testicular cells to recreate the native architecture of testicular tissue found *in vivo*. Recent advances in 3-D bioprinting can place cells into geometries at fine resolutions comparable to microarchitectures found in native tissues, and therefore hold promise as a tool for the development of a biomimetic *in vitro* system for human spermatogenesis. This study assessed the utility of bioprinting technology to recreate the precise architecture of testicular tissue and corresponding spermatogenesis for the first time. We printed testicular cell-laden hollow microtubules at similar resolutions to seminiferous tubules, and compared the results to testicular organoids. We show that the human testicular cells retain their viability and functionality post-printing, and illustrate an intrinsic ability to reorganize into their native cytoarchitecture. This study provides a proof of concept for the use of 3-D bioprinting technology as a tool to create biomimetic human testicular tissues.

## Introduction

### Need for Regenerative Male Fertility Therapies

Survival rates for pediatric cancer patients are steadily rising due to improvements in early detection and treatments, with recent estimates at 84%.(1–3) The increased survival rate has brought about considerations for quality of life, and in particular fertility restoration because of the gonadotoxic side effects of cancer treatments.(4) Pre-pubertal boys may cryopreserve testis tissue in hopes that a future safe regenerative treatment will be developed to restore fertility, and research amongst this population has demonstrated that 75% of childhood cancer survivors want future offspring.(5)

### Experimental options for fertility preservation

Potential experimental treatments to restore fertility using these prepubertal tissues include their use for autologous transplantation, and the isolation and *in vitro* expansion of spermatogonial stem cells (SSCs) - the stem cell precursors to spermatozoa – for autologous transplantation.(6, 7) However, these strategies present the risk of re-introducing malignant cancer cells to the post-treatment patient, and therefore safety issues must be addressed before human trials can take place.(8, 9) A safer alternative being explored is *in vitro* spermatogenesis, wherein cryopreserved prepubertal tissue is matured in an *in vitro* setting.(10) Spermatozoa from these systems can be isolated and used for *in vitro* fertilization (IVF).

Although *in vitro* systems can generate spermatozoa from animal SSCs, these techniques have not translated successfully to human cells.(9, 11–13) Of the successful animal systems, organotype culture was the first to generate spermatozoa,(14) presumably due to its preservation of the testicular cytoarchitecture. Indeed, the cytoarchitecture of testicular tissue is one of the most complex in the body, and its improper establishment is correlated with impaired spermatogenesis.(11)

### 3D-bioprinting as a tool for generating biomimetic testicular tissue *in vitro*

3D-bioprinting has emerged as a useful tool for fabricating biomimetic living tissues by the automated layer-by-layer assembly of cells. This technology can build tissues directly into culture plates with geometries at micrometer resolutions similar to native tissues.(15) Hollow structures can be generated by printing sacrificial materials into these spaces to be removed post-printing, and coaxial nozzle printheads can exploit this principle to fabricate hollow tubule structures.

The purpose of this study was to assess their feasibility and performance of bioprinted human adult testicular cells compared to current gold standard organoids as a first step towards the goal of regenerating human testicular tissue *in vitro*. We used a bioprinter whose microfluidic technology enables precise patterning of functional biomaterials to recreate microtubules resembling the size and shape of native seminiferous tubules – the niche where spermatogenesis takes place, and we analyzed their viability, gene expression, and cytoarchitecture after 12 days in culture. This study serves as a proof of principle for 3-D bioprinting testicular tissues, and offers recommendations for their further development into systems for human *in vitro* spermatogenesis.

## Materials and methods

### Ethics approval

Ethics approval for the use of human tissues was obtained through the University of British Columbia Clinical Research Ethics Board (CREB approved protocol H18-03543). All participants gave informed, written consent as per Protocol 1.6, and tissue was obtained as per Protocol 1.3.

### Tissue dissociation and cell culture

A testicular biopsy was transported to the lab on ice in Hypothermosol® FRS (STEMCELL Technologies, 07935), and digested as previously described.(16) Cells were sorted into somatic and germ fractions by overnight plating: the adherent cells were expanded in a 1:1:1 mixture of Endothelial Cell Growth Medium (Promocell, C-22010), Sertoli Cell Medium (Sciencell, 4521), and Leydig Cell Medium (Sciencell, 4511), and non-adherent cells were transferred the next day to a CellAdhere™ Laminin 521 (STEMCELL Technologies, 77004)-coated plate, and expanded in SSC expansion media as previously described.(17)

### 3-D bioprinting

Bioprinting was accomplished using an RX1™ bioprinter (Aspect Biosystems™) and a CENTRA™ coaxial microfluidic printhead (Aspect Biosystems™). Aspect Studio software was used to generate a ring design, since when printing directly into a solution, moving the printhead in a ring shape prevents clumping. Primary germ and somatic cells were mixed together at a ratio of 1:4 into AGC-10 matrix™ (Aspect Biosystems™) to a final concentration of 35 million cells/mL, and printed with 6% polyvinyl alcohol (PVA, Aspect Biosystems™) as the sacrificial core material. CAT-2™ crosslinker (Aspect Biosystems™) was used. Prints were cultured in Ultra-Low Adherent Plates for Suspension Culture (STEMCELL Technologies, 38071) for 12 days in SSC expansion media with the following additions: 100 ng/mL follicle stimulating hormone from human pituitary (FSH, Millipore Sigma, F4021), 10 ng/mL luteinizing hormone from human pituitary (LH, Sigma, L6420), 1 μM metribolone (Toronto Research Chemicals, M338820), 100 ng/mL human recombinant Bone Morphogenic Protein 4 (BMP4, Peprotech, AF-120-05ET), 100 ng/mL animal-free recombinant human Stem Cell Factor (SCF, Peprotech, AF-300-07), and 10 μM all-trans retinoic acid (RA, STEMCELL Technologies, 72262), and the removal of Glial Cell Derived Neurotrophic Factor (GDNF).

### Organoid generation

Testicular organoids were made by mixing germ and somatic cells at a 1:4 ratio and placing them into AggreWell800™ microwells (STEMCELL Technologies, 34811) overnight, followed by culture under the same conditions as the prints.

### Reverse Transcription Qualitative Polymerase Chain Reaction (RT-qPCR)

RNA was extracted with an RNeasy Plus Micro Kit (Qiagen, 74034), and checked for integrity using an Agilent 2200 Tapestation System and High Sensitivity RNA Screentape system (Agilent, 5067-5579-81). cDNA was generated using iScript Reverse Transcription Supermix (Bio-Rad, 1708840), followed by pre-amplification with SsoAdvanced™ PreAmp Supermix (Bio-Rad, 1725160) with a Tetrad2 Peltier Thermal Cycler (Bio-Rad). RT-qPCR was done with SsoAdvanced™ Universal SYBR® Green Supermix (BioRad, 1725270) on a LightCycler96 (Roche). Pre-amplification primers and regular primers used were PrimePCR™ SYBR® Green Assays as listed in **Table 1.** Analyses was done in Excel and GraphPad Prism Software. Ct values were normalized to Glyceraldehyde 3-Phosphate Dehydrogenase (GAPDH). Technical replicate outliers were detected using Grubbs’ Test, with α=0.05. Results of RT-qPCR are presented as the average Relative Quantification (RQ=2^−ΔΔCt^) values and standard deviations of the biological replicates. Any undetected samples were given a Ct value of the maximum detectable cycles.

**Table 1.**
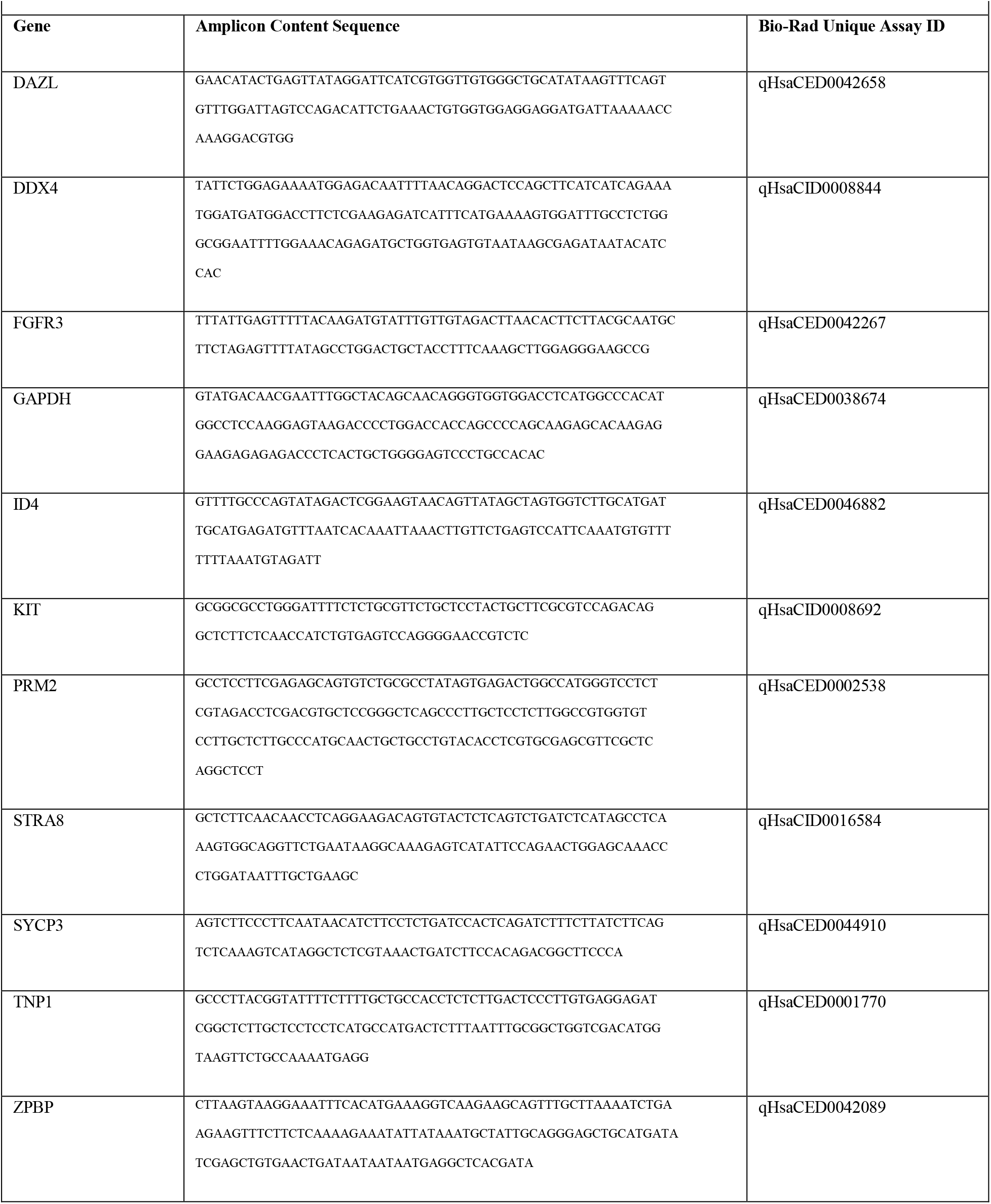
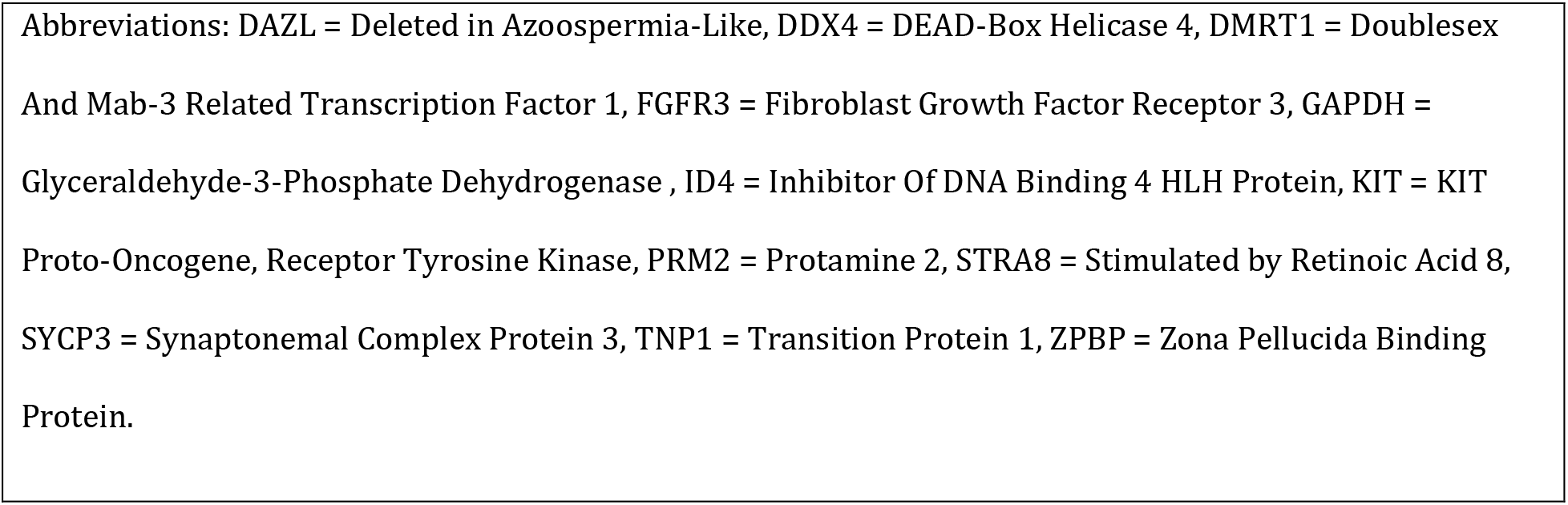
Amplicon content sequences (amplicon sequence with additional base pairs added to the beginning and/or end of the sequence), and Unique Assay IDs for Bio-Rad PrimePCR™ primer pairs used in RT-qPCR.

### Viability assay

Viability of the bioprinted fibers was assessed after 24 hours using a LIVE/DEAD™ Sperm Viability Kit (Invitrogen, L7011) as per the manufacturer’s instructions, and imaged using a Zeiss AxioObserver confocal microscope. Viability was quantified using ImageJ software by making each separate color channel image into an 8-bit binary image, and then thresholding it to mask the cells, followed by segmenting of merged cells using the watershedding command, and counting using the Analyse Particles command. Results are presented as the average and standard deviation of three biological replicates.

### Immunocytochemistry

Cells were fixed for 15 minutes in 4% paraformaldehyde solution (PFA, Thermo Scientific, J19943-K2), permeabilized for 15 minutes in 0.1% Triton X-100 (Sigma, X100) in phosphate buffered saline (PBS), and blocked for 2 hours in 5% normal goat serum (NGS, Abcam, ab7481) in PBS. Primary antibodies were diluted in PBS as follows: anti-Hydroxy-Delta-5-Steroid Dehydrogenase, 3 Beta- And Steroid Delta-Isomerase 1 (HSD3β, Novus Biologicals, NB110-7844) 1:200, anti-Myosin Heavy Chain 11 (MYH11, Abcam, ab212660) 1:1000, anti-SRY-Box Transcription Factor 9 (SOX9, Abcam, ab76997) 1:500, anti-Thy-1 Cell Surface Antigen (THY1/CD90, Abcam, ab133350) 1:200, 1:300, anti-Insulin Like 3 (INSL3, Novus Biologicals, NBP1-81223) 1:500, anti-CD34 Molecule (CD34, Abcam, ab81289) 1:250, anti-GDNF Family Receptor Alpha 1 (GFRA1, Abcam, ab84106) 1:200, anti-G-Protein Coupled Receptor 125 (GPR125, Abcam, ab51705) 1:200, anti-Stage-specific embryonic antigen-4 (SSEA4, Abcam ab16287) 1:300, anti-STRA8 (Millipore ABN1656), and anti-Actin Alpha 2, Smooth Muscle (ACTA2, Thermofisher, 14-9760-82) 1:500, and incubated overnight at 4°C in the dark. Cells were rinsed 3 times with PBS for 15 minutes each. Goat anti-Rabbit IgG (H+L) Highly Cross-Adsorbed Secondary Antibody Alexa Fluor 488 (Thermofisher, A-11034) or Goat anti-Mouse IgG (H+L) Highly Cross-Adsorbed Secondary Antibody Alexa Fluor 568 (Thermofisher, A-11031) were diluted 1:200 in PBS and incubated with the cells for 4 hours at 4°C in the dark, followed by 3 rinses in PBS for 15 minutes each. 4′,6-diamidino-2-phenylindole (DAPI, Abcam, ab228549) was diluted to 2.5 μM in PBS and added to the cells for 15 minutes in the dark. Cells were imaged using a Zeiss AXio Observer microscope, and images were processed with ZEN Blue and ImageJ software.

### Immunofluorescence staining of the 3-D organoids and prints

Embedding, sectioning and immunofluorescent staining of the organoids was done as previously described.(18) Embedding, sectioning and staining of the prints was done the same way with the following modifications: prints were fixed in 4% PFA solution saturated with calcium chloride (Sigma, C4901) to prevent dissolving of the bioprinted matrix, and after sectioning fixed a second time in 4% PFA for 30 minutes followed by permeabilization with 0.1% Triton-X100 in PBS. Whenever 0.1% Tween is used for the organoids, it is replaced by PBS for the prints to avoid washing away the cells. Primary antibodies were diluted as follows: 1:1000 anti-SOX9, 1:500 anti-ACTA2, 1:500 anti-INSL3, 1:300 anti-SYCP3 (Novus Biologicals, NB300-232). Secondary antibodies were Goat anti-Mouse IgG (H+L) Cross-Adsorbed Secondary Antibody Alexa Fluor 647 (Thermofisher, A-21235) or Goat anti-Rabbit IgG (H+L) Cross-Adsorbed Secondary Antibody Alexa Fluor 647 (Thermofisher, A27040), diluted 1:2000. Sections were imaged using a Zeiss AXio Observer microscope, and images were processed using ZEN Blue and ImageJ software.

### Hematoxylin and Eosin (H&E) staining

Embedding and sectioning was completed as described in Immunofluorescence staining. Organoid sections were incubated with 0.1% Tween 20 in PBS for 10 minutes at 37⁰C, while the print sections were not. The slides were dipped as follows: hematoxylin solution (Sigma Aldrich, MHS32)-3 minutes, tap water-1 minute x 2, 0.1% sodium bicarbonate (Sigma S5761) solution-1 minute, tap water-1 minute, 95% ethanol-1 minute, Eosin Y reagent (Sigma Aldrich, HT110216)-1 minute, 95% ethanol-1 minute, 100% ethanol-1 x 2, xylene (Fisher Scientific X5-500)-3 minutes. 1-2 drops of CytoSeal (Thermofisher, 8312-4) were placed on each before mounting a coverslip. Organoids were imaged using an Olympus BX-UCB (PerkinElmer Life Sciences) and VECTRA software, while prints were imaged using an EVOS XL Core imaging System (Invitrogen).

### Statistics

Statistics were performed using GraphPad Prism software. Each experiment was performed in biological triplicate. Significance was determined by comparing ΔCt values using a student’s unpaired two-tailed t-test with α=0.05.

## Results

### Establishment of testicular cell cultures

Primary testis cells were isolated from a human testicular biopsy, separated by differential plating into germ and somatic fractions, and expanded *in vitro* (**Figure 1A**). The main somatic cells in the testicular niche are: Sertoli cells, the master regulators of spermatogenesis and main component of seminiferous tubules; Leydig cells, an interstitial cell that produces the testosterone necessary for Sertoli cell function; peritubular myoid cells, a smooth muscle cell type that confers contractility to the seminiferous tubules in order to advance spermatozoa towards the epididymis; and endothelial cells, vascular cells which secrete factors important to the morphogenesis and homeostasis of the niche.(19) Immunocytochemistry showed that somatic cultures were largely positive for SOX9^+^ Sertoli cells (SCs)(20), with smaller pockets of MYH11^+^ peritubular myoid cells (PTMs)(21) and CD34^+^ endothelial cell progenitors (ECs) (**Figure 1C**).(22) Expression of Leydig cell markers HSD3β(23) and INSL3(23) were absent, indicating either their loss or de-differentiation (**Figure 1C**). The proliferative state noted in the Sertoli cells was also suggestive of de-differentiation, in keeping with a previous report of adult human Sertoli cell culture.(24) Immunocytochemistry showed homogeneous expression of SSC genes CD90, GFRA1, SSEA4, GPR125, and STRA8 (**Figure 1B**),(25–28) while expression of somatic cell markers ACTA2(27) and SOX9 were negative (**Figure 1B**). Since they were otherwise presenting an undifferentiated SSC phenotype, their expression of STRA8, a retinoic acid-stimulated gene required for entry into meiosis, was an interesting observation. One explanation is that it was induced by unnatural *in vitro* culture conditions. This is consistent with studies showing that SSCs lose their resistance to retinoic acid stimulation when removed from their 3-D microenvironment and upregulate STRA8.(29, 30)

**Figure 1.**
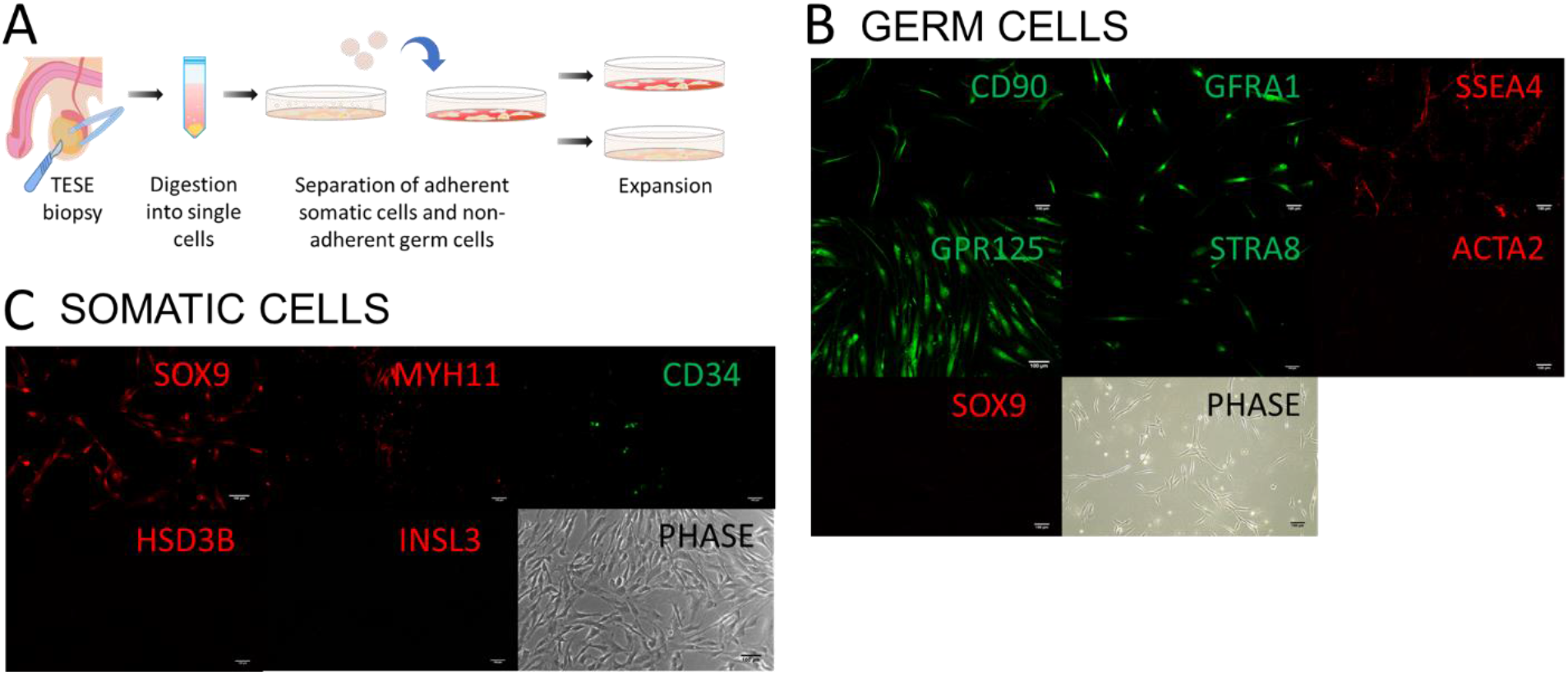
Primary testicular cell isolation, expansion and characterization. **A)** Schematic representing the isolation and separation of testicular cells into germ and somatic fractions: a testicular biopsy was dissociated into single cells, plated overnight, and non-adherent germ cells were collected and plated on a laminin-coated plate. **B)** Immunocytochemistry staining of the germ cells for spermatogonial stem cell markers CD90, GFRA1, SSEA4, GPR125, and STRA8, and the testicular somatic cell markers ACTA2 and SOX9. **C)** Immunocytochemistry staining of the somatic cells for the Sertoli cell marker SOX9, the peritubular myoid cell marker MYH11, the endothelial progenitor cell marker CD34, and the Leydig cell markers HSD3B and INSL3. All scale bars are 100 μm. **Abbreviations:** TESE = testicular sperm extraction, FGF2 = Fibroblast Growth Factor 2, LIF = Leukemia Inhibitory Factor, EGF = Epidermal Growth Factor, GDNF = Glial Cell-Derived Growth Factor, SOX9 – SRY-Box Transcription Factor 9, MYH11 = Myosin Heavy Chain 11, CD34 = CD34 Molecule, HSD3B = Hydroxy-Delta-5-Steroid Dehydrogenase, 3 Beta- And Steroid Delta-Isomerase 1, INSL3 = Insulin-Like Protein 3, CD90 = Thy1 Cell Surface Antigen, GFRA1 = GDNF Family Receptor Alpha 1, SSEA4 = Stage-Specific Embryonic Antigen-4, GPR125 = G-Protein Coupled Receptor 125, STRA8 = Stimulated by Retinoic Acid 8, ACTA2 = Actin Alpha 2, Smooth Muscle.

### Generation of 3-D bioprinted testicular tubules

Expanded primary testicular cells were either bioprinted directly into wells containing media or seeded into microwells to form organoids as a control (**Figure 2A,C**). The microfluidic printhead used incorporates separate channels for crosslinker, buffer, core, and shell materials (**Figure 2A**). To print, testicular cell-laden bioink was loaded into the shell channel, and the sacrificial material PVA was loaded into the core channel. As the core channel material flows into the printhead it is surrounded by the shell channel material to produce concentric streamlines by coaxial flow focusing. The crosslinker then enters the channel, surrounding the core-shell material, and as the fiber exits the printhead, the bioink rapidly crosslinks and a hollow cell-laden tubule is formed. Tubule diameter was fine-tuned by adjusting the pressures of the shell, core and crosslinker channels, and the speed of extrusion. We found that shell, core and crosslinker pressures of 150 mbar, 300 mbar, and 125 mbar, extruded at a speed of 25 mm/s, produced a tubule diameter of ~500-600 μm (**Figure 2A**). Tubules were deposited directly into media and the printhead was programmed to move around the well in a circular motion to prevent clumping. Each print was the length of 5 complete circles, or roughly 20-30 cm. Prints were found to be on average 93.4±2.4% viable after 24 hours by LIVE/DEAD™ staining (**Figure 2B**), confirming that the cells were not damaged during the bioprinting process. Organoid controls were generated in parallel by combining germ and somatic cells 1:4 into AggreWell800™ microwells for 24 hours (**Figure 2C**).

**Figure 2.**
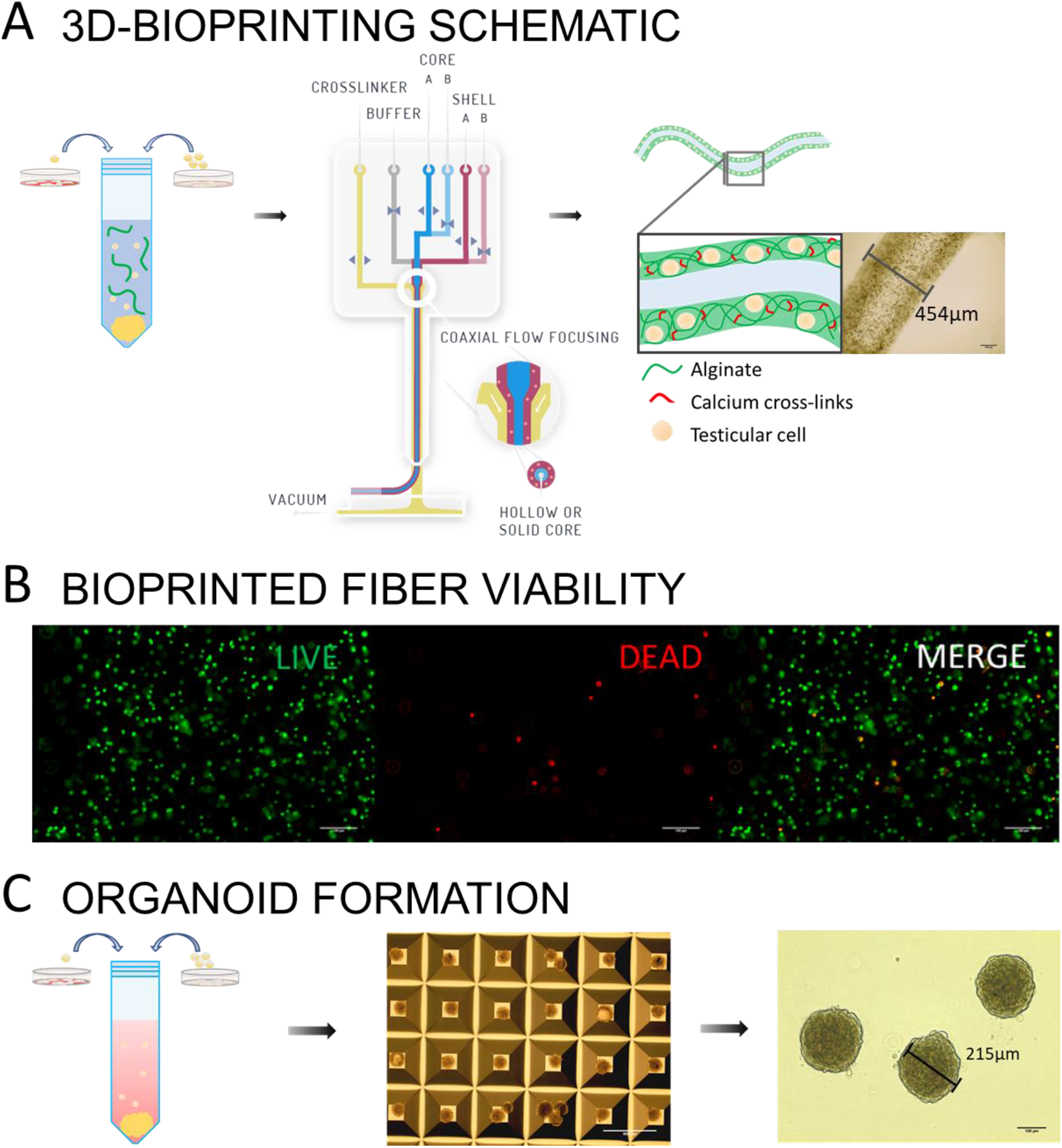
3D-bioprinting schematic and viability of 3D-printed cells. **A)** Schematic representing the 3D-bioprinting process to generate hollow testicular cell-laden tubules: germ and somatic cells were combined 1:4 into an un-crosslinked alginate-based bioink and extruded through a microfluidic printhead which incorporates separate microfluidic channels for crosslinker, buffer, core and shell, and uses coaxial flow focusing to create core-shell fibers. **B)** Viability staining after 24 hours. Quantification using image processing software was used to calculate an average viability of 93.4±2.4%. **C)** Organoid formation: germ and somatic cells were combined 1:4 into AggreWell800™ microwells for 24 hours. All scale bars are 100 μm.

### Characterization of 3-D bioprinted testicular tubules

Prints and organoids were cultured in media supplemented with hormones FSH, LH, and androgen to promote somatic cell function,(31) and growth factors LIF, EGF, SCF, BMP4, and retinoic acid to promote germ cell differentiation and survival (**Figure 3A**).(32–40). After 12 days we analyzed their morphology and gene expression by H&E staining, immunofluorescence, and RT-qPCR assays.

**Figure 3:**
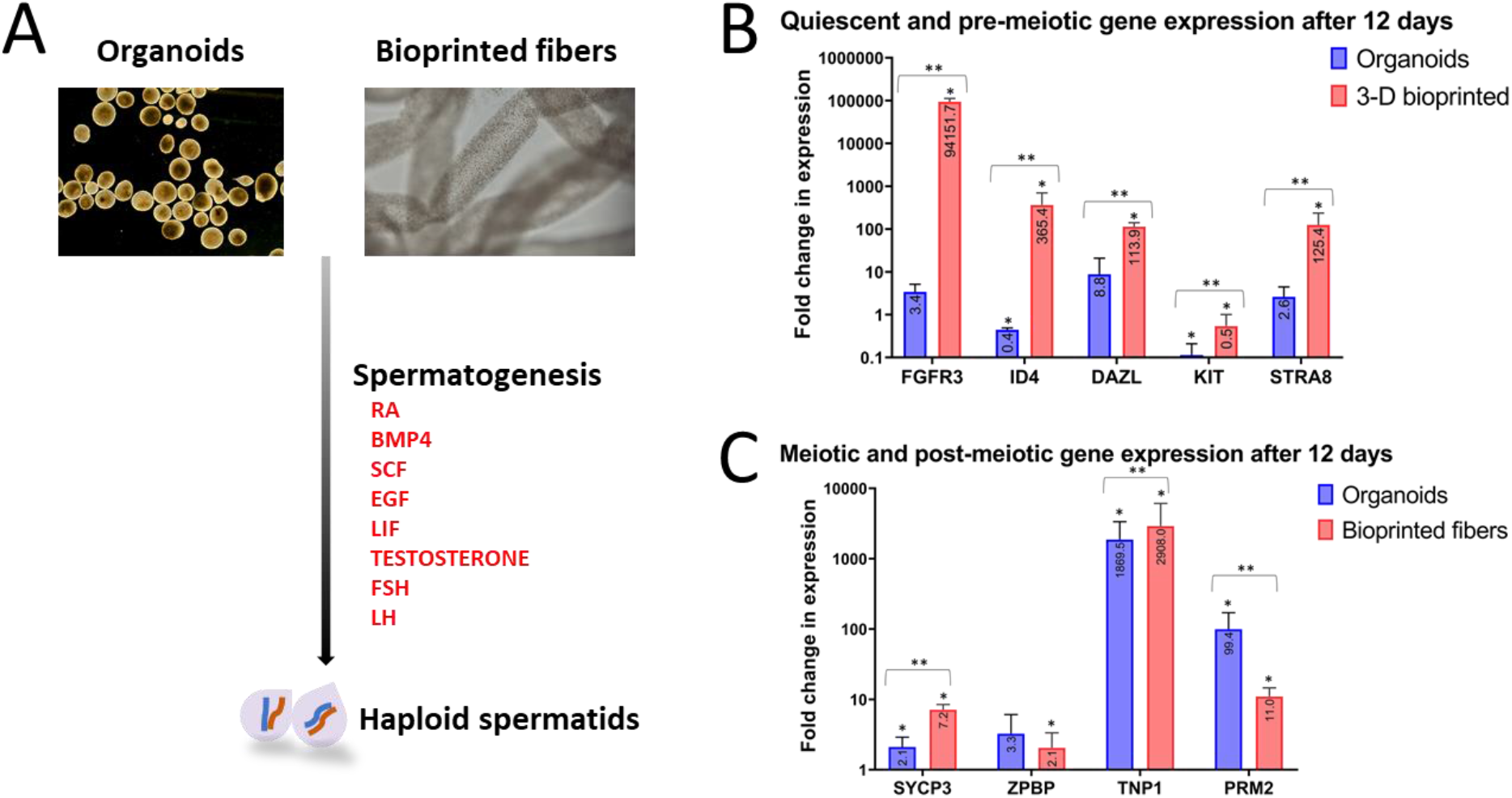
Generation and functional gene expression analyses of the prints and organoids. **A)** Schematic representing the generation and culture of the prints and organoids. **B)** Fold change in gene expression for the quiescent spermatogonial stem cell genes FGFR3 and ID4, the spermatogenesis regulatory gene DAZL, and the pre-meiotic genes KIT and STRA8. **C)** Fold change in gene expression for the meiotic gene SYCP3, and the post-meiotic spermiogenesis genes ZPBP, TNP1 and PRM2. * indicates a statistically significant difference from day 0 cultures, and ** indicates a statistically significant difference between prints and organoid controls. **Abbreviations:** RA = retinoic acid, BMP4 = Bone Morphogenic Protein 4, SCF = Stem Cell Factor, EGF = Epidermal Growth Factor, LIF = Leukemia Inhibitory Factor, FSH = follicle stimulating hormone, LH = luteinizing hormone, SSC = spermatogonial stem cell, FGFR3 = Fibroblast Growth Factor 3, ID4 = Inhibitor of DNA Binding 4, HLH Protein, DAZL = Deleted in Azoospermia-Like, KIT = KIT Proto-Oncogen, Receptor Tyrosine Kinase, STRA8 = Stimulated by Retinoic Acid 8, SYCP3 = Synaptonemal Complex Protein 3, ZPBP = Zona Pellucida Binding Protein, TNP1 = Transition Protein 1, PRM2 = Protamine 2.

### Functional gene expression

During the cycle of spermatogenesis, the balance of SSC self-renewal and differentiation is critical to the continuous regeneration of spermatozoa, and this balance relies upon the organization and architecture of the testicular niche.(41) Therefore, to assess the effects of the bioprinted architecture on this balance we analyzed the prints and organoid controls for changes in gene expression for self-renewing SSC markers ID4(42) and FGFR3,(43) and differentiating markers KIT(44) and STRA8 (**Figure 3B**).(45) Interestingly, prints were noted to upregulate ID4 and FGFR3 expression considerably [+365-fold and +94,152-fold], whereas organoids downregulated ID4 expression [-2-fold] and did not significantly change FGFR3 expression (**Figure 3B**), perhaps due to increased cell contact inhibition caused by their compactness compared to the prints. We noted a pattern of KIT downregulation and STRA8 upregulation in both the prints and organoids [prints: -2-fold, 125-fold; organoids: −11-fold, +3-fold] (**Figure 3B**). In accordance with the increase in expression of these spermatogenic genes, expression of DAZL, a master translational regulator required for SSC self-renewal and differentiation(46, 47) was also upregulated in both the prints and organoids [114-fold and 9-fold] (**Figure 3B**).

During spermatogenesis, SSCs undergo a division process specific to germ cells called meiosis to produce haploid daughter cells, each containing unique samples of the parent cell’s genetic material. Haploid cells then undergo a morphological transformation to become spermatozoa during a maturation phase called spermiogenesis.(48) To assess the effects of the printed hollow tubule structure on meiosis and spermiogenesis, we analyzed changes in gene expression for SYCP3, a marker of meiosis;(49) ZPBP, a gene required for the development of a specialized egg-penetrating organelle called an acrosome;(27, 50) and TNP1 and PRM2, two genes which encode proteins that replace histones during chromosome compaction.(27) Both the prints and organoids upregulated SYCP3 expression [7-fold and 2-fold] (**Figure 3C**). Conversely, we found that in the prints ZPBP expression was upregulated [2-fold], while it was not significantly changed in the organoids (**Figure 3C**). Both TNP1 expression and PRM2 expression in the prints and organoids were substantially upregulated [prints: 2,908-fold, 11-fold, organoids:1,870-fold, 99-fold] (**Figure 3C**).

### Cytoarchitecture and morphology

Prints and organoid controls were fixed, embedded and stained for nuclei and cytoplasm contents to assess any tissue-like organization. The prints established tubular structures within some sections of bioprinted tubules, illustrating a strong intrinsic ability to self-organize (**Figure 4A**). Immunofluorescence showed that SOX9+ Sertoli cells and SYCP3^+^ meiotic germ cells localized towards the luminal areas of these structures, while ACTA2^+^ myoid cells localized peripheral to Sertoli cells, and INSL3^+^ Leydig cells clustered together synonymous with human *in vivo* cytoarchitecture (**Figure 4A**). The presence of INSL3^+^ Leydig cells confirmed their de-differentiated status in 2-D culture. Organoids similarly developed tubular inner structures (**Figure 4B**) and immunofluorescence showed that SOX9^+^ Sertoli cells co-localized with SYCP3^+^ germ cells in the tubular structures, while ACTA2^+^ peritubular myoid cells and INSL3^+^ Leydig cells appeared to be localized peripheral to these structures (**Figure 4B**).

**Figure 4:**
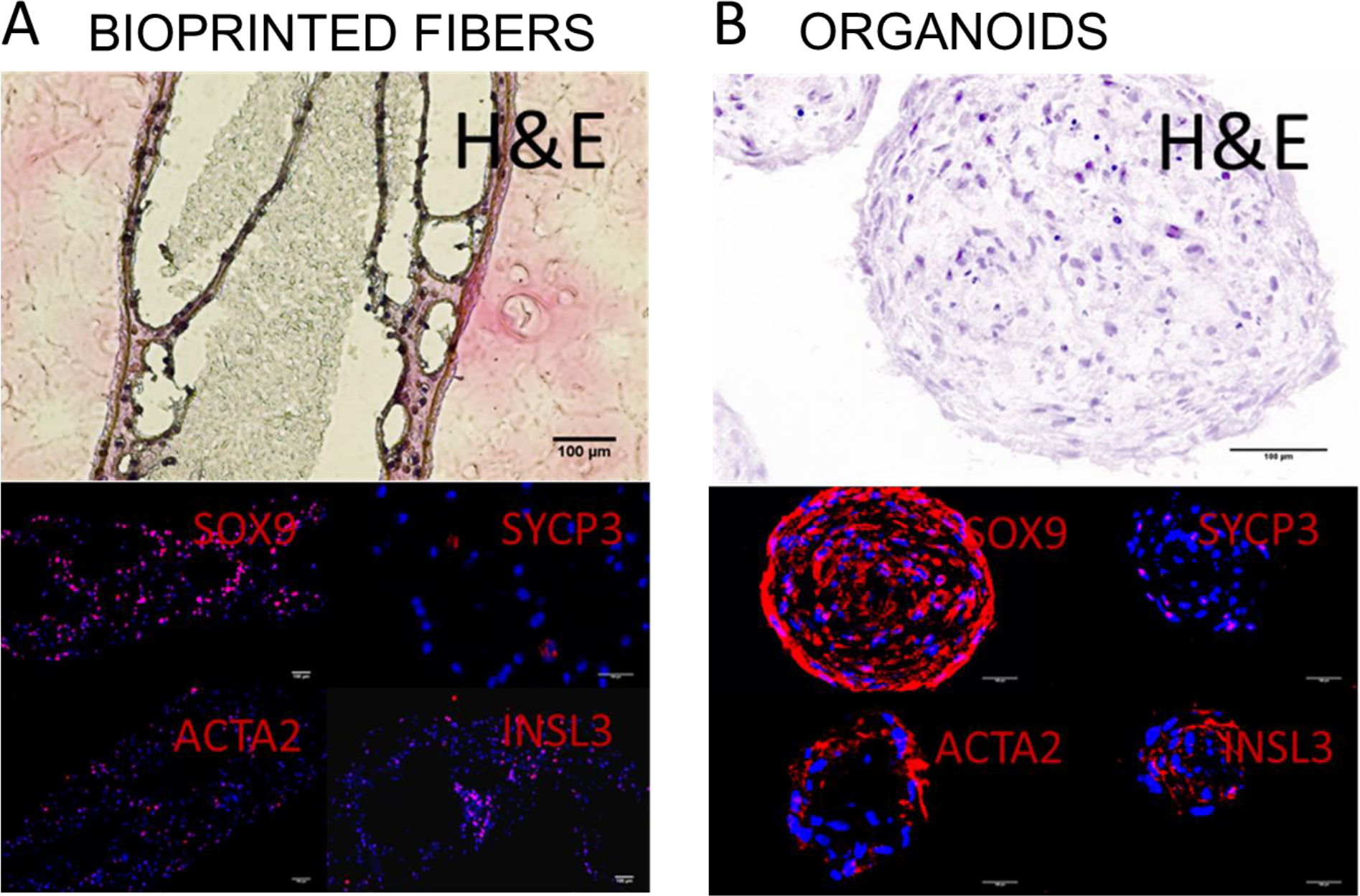
Morphology of the testicular prints and organoids. **A)** H&E staining of the prints and immunocytochemistry staining for the Sertoli cell marker SOX9, the meiotic germ cell marker SYCP3, the peritubular myoid cell marker ACTA2, and the Leydig cell marker INSL3. The dashed white lines show the outlines of the tubule sections. **B)** H&E staining of the organoids and immunocytochemistry staining for the Sertoli cell marker SOX9, the meiotic germ cell marker SYCP3, the peritubular myoid cell marker ACTA2, and the Leydig cell marker INSL3. All scale bars are 100 μm. **Abbreviations:** H&E = Hematoxylin and Eosin, SOX9 =SRY-Box Transcription Factor 9, SYCP3 = Synaptonemal Complex Protein 3, ACTA2 = Actin Alpha 2, Smooth Muscle, INSL3 = Insulin Like 3.

## Discussion

This study investigated the utility of a novel bioprinting strategy to recreate biomimetic human testicular tissues with regenerative potential. We used a microfluidic coaxial printhead design to bioprint microtubules similar in size to the seminiferous tubules which house the spermatogenic niche using human testicular cells, and we compared their resulting cytoarchitecture and functional gene expression to self-assembled organoid controls.

An important finding was the high viability (93.4±2.4%) of the testicular cells post-printing. Extrusion bioprinting imparts shear stress on cells, which can result in poor viability.(51) The use of microfluidic bioprinting solves this problem by extruding cell suspensions in un-crosslinked materials separately from a crosslinking agent, such that gelation of the bioink occurs post-extrusion, greatly reducing shear stress to enable high viability of even sensitive cells such as stem cells.(52)

A second important finding was the re-establishment of an *in vivo*-like cytoarchitecture. The correct organization of human testicular cells in 3-D systems has been limited to date, even with the use of decellularized testicular ECM as a natural biomimetic scaffold.(53) We noted the development of tubular inner structures of Sertoli cells and germ cells in both the prints and organoid controls, and the peripheral localization of Leydig and peritubular myoid cells in accordance with *in vivo*-like organization. An explanation for this difference compared to other human testicular organoid studies may be the noted de-differentiation of the testicular cells in 2-D culture, leading to progenitor-like morphogenic behavior. This correlates with another study wherein adult human testicular cells plated in 2-D, in the presence of the testicular morphogenic growth factors FGF9 and FGF2,(54) aggregated into organized 3-D structures resembling *in vivo* like cytoarchitectures.(55) A second contribution to the organization of the testicular cells in this study may have been their noted maturation following culture in the 3-D bioprinted and organoid systems. When human testicular organoids are made from pre-pubertal cells in the absence of hormonal stimulation they have been found to organize only partially, in an inside-out manner where the tubular compartment is localized to the outer edge and the interstitial compartment is internalized.(29) Furthermore, when testicular organoids are made from immortalized testicular cells they are unable to organize,(56) suggesting that proliferative arrest, a property of maturation, may be critical to allowing organization to take place. Nevertheless, while the prints in our study developed tubular and interstitial organization similar to *in vivo* cytoarchitecture, these structures lacked *in vivo*-like scale, and by association, complexity, illustrating the need for precise placement of interstitial, peritubular and tubular cells into defined spatial compartments. To this end, microfluidic printheads incorporating multiplex coaxial flow focusing may be useful tools in future bioprinted designs to create multicompartmental core-shell structures.

Finally, assessing the functionality of the prints was key to interpreting their utility for *in vitro* regeneration of spermatozoa. The ability of the prints to upregulate functional genes corresponding to SSC self-renewal, meiosis and spermiogenesis, on par with the organoid system, confirmed that testicular cells retain their functionality post-printing. Furthermore, the prints were noted to express substantially greater levels of genes associated with SSC self-renewal and differentiation compared to organoid controls, denoting an improvement in the regulatory niche within the bioprinted architecture.

## Conclusions and future directions

This study demonstrates for the first time that bioprinting of adult human testicular cells provides a feasible platform for potential regenerative therapies through *in vitro* spermatogenesis. Our findings demonstrate that bioprinted cells generate biomimetic cytoarchitecture, while retaining cell viability and re-establishing a functional spermatogenic niche. Compared to organoid controls, bioprinted tubules better supported quiescent SSC populations important for sustaining long term-culture and spermatogenesis, and demonstrated upregulation of both meiotic and post-meiotic gene expression both important for use as a future regenerative platform. Future studies will be necessary to further engineer the tubular architecture and establish more efficient luminal and interstitial culturing systems.

## Acknowledgements

The authors would like to acknowledge the Vancouver Prostate Centre for funding support, and Aspect Biosystems for technical support in bioprinting. We would also like to acknowledge the Histology Core and the Lange Lab at the Vancouver Prostate Centre for their generous assistance in staining and imaging the organoids and 3-D bioprinted fibers.

## Conflicts of interest

The authors have a patent pending for the biomimetic seminiferous tubule model.

## Data availability statement

The data that support the findings of this study are available upon reasonable request from the authors.

